# Hypothetical LOC Genes as Biomarkers of Spaceflight Adaptation: A Comparative Study from ISS, Suborbital, and Earth-Based Experiments

**DOI:** 10.1101/2025.09.12.675876

**Authors:** Özge Demir, Ebru Çam, Batuhan Yolver, Kürşat Kolay, Merve Reyhan Sakar, Beyza Aydın, Cihan Taştan

## Abstract

Microgravity constitutes one of the most profound environmental stressors encountered by humans during spaceflight, capable of altering fundamental cellular processes and gene regulatory networks. While the effects of spaceflight on well-characterized protein-coding genes have been widely documented, little is known about the behavior of uncharacterized or poorly annotated genomic regions under these conditions. LOC (Locus) genes, often classified as long non-coding RNAs and excluded from conventional analyses, represent a largely unexplored component of the human transcriptome. In this study, we systematically investigated the transcriptional responses of LOC genes as part of the MESSAGE (Microgravity Associated Genetics) Science Mission, Türkiye’s first human space biology initiative. Peripheral blood samples were collected from astronauts across five mission phases: pre-launch baseline, post-suborbital flight (∼100 km), and on International Space Station (ISS) Days 4, 7, and 10 (∼400 km). RNA-Seq analyses revealed six LOC genes with statistically significant expression changes (p < 0.05, Kruskal–Wallis test), alongside additional transcripts that, while not statistically significant, exhibited biologically meaningful temporal fluctuations. These dynamic profiles included continuous upregulation, transient activation with subsequent return to baseline, and delayed induction at later ISS stages, highlighting the functional diversity of LOC responses. To assess their translational potential, Open Reading Frame (ORF) analyses were performed on significant transcripts, revealing conserved ORF structures—most notably identical ORF33 sequences in LOC124905103 and LOC124900480— suggesting coding capacity. Phylogenetic analyses further supported evolutionary clustering consistent with expression and ORF similarities. Collectively, these findings challenge the notion of LOC genes as transcriptional noise, instead positioning them as candidate biomarkers and functional elements of microgravity adaptation. By extending space biology research into the “dark genome,” this study provides novel insights with potential implications for astronaut health monitoring and therapeutic development in long-duration missions.

## 1. Introduction

Spaceflight introduces a novel set of environmental stressors, with microgravity standing out as a key factor that profoundly affects human physiology at the cellular and molecular levels. Gravity, a fundamental physical constant on Earth, has guided the evolution of all biological systems. It is now well established that gravitational unloading impacts mechanotransduction pathways that govern cytoskeletal organization, signal transduction, cell cycle regulation, metabolic homeostasis, and gene expression [1, 2, 3]. Disruptions in these fundamental processes under microgravity represent a significant focus in the field of space biology. Recent increases in orbital and suborbital missions, such as those conducted on the International Space Station (ISS) (∼400 km) and during short-duration suborbital flights (∼100 km), have made it possible to investigate how microgravity alters immune function, oxidative stress, mitochondrial integrity, and transcriptional activity [5, 4, 7, 6]. Studies have demonstrated that microgravity can suppress immune activation, destabilize cytoskeletal structures, and increase the generation of reactive oxygen species (ROS), which in turn causes molecular damage to DNA, lipids, and proteins [8, 9]. While previous transcriptomic studies have largely focused on well-characterized, protein-coding genes, much less is known about the behavior of unannotated or poorly characterized genomic regions under microgravity. LOC (Locus) genes—transcripts labeled as “uncharacterized” due to incomplete functional annotation—fall into this underexplored category. Many LOC genes are classified as long non-coding RNAs (lncRNAs), exhibit low expression levels, and are often excluded from conventional analysis pipelines due to technical challenges [10]. To date, no comprehensive study has systematically assessed the expression profiles of LOC genes in the context of microgravity exposure (as of June 2025). This study, conducted as part of Türkiye’s first human spaceflight program under the MESSAGE (Microgravity Associated Genetics) Science Mission, addresses this critical knowledge gap. Peripheral blood samples were collected from astronauts at five time points: prior to launch, following a suborbital flight, and during ISS mission days 4, 7, and 10. RNA-Seq analyses were performed to examine temporal gene expression patterns under microgravity conditions. Statistical evaluation of expression changes was conducted using the non-parametric Kruskal-Wallis test to accommodate a small sample size and non-normal distribution [15]. This study, conducted under Türkiye’s first human spaceflight program within the MESSAGE Science Mission, was designed to address this knowledge gap by systematically characterizing LOC gene expression under microgravity. Peripheral blood samples were collected from astronauts at five mission phases, spanning pre-launch baseline, post-suborbital flight, and three ISS time points (Days 4, 7, and 10). RNA-Seq analyses were performed to capture time-resolved transcriptional dynamics, with statistical evaluation carried out using the Kruskal–Wallis test to account for limited sample size and non-normal distributions. By integrating expression profiles with phylogenetic and ORF analyses, this study aimed to determine whether LOC genes— traditionally regarded as uncharacterized or non-coding—exhibit functional relevance in spaceflight. We hypothesized that these transcripts may serve as regulatory elements or encode peptides with adaptive significance and thus represent previously overlooked biomarkers of human physiological response to microgravity.

## 2. Material and Methods

### Sample Collection and Study Context

This study was conducted as part of Turkey’s first human spaceflight mission, the MESSAGE (Microgravity Associated Genetics) Science Mission, coordinated by TÜBİTAK Space Technologies Research Institute and the Turkish Space Agency (TUA). Peripheral blood samples were collected from three astronauts who participated in an orbital mission aboard the International Space Station (ISS) and one astronaut who completed a suborbital flight at an altitude of approximately 100 km. Samples were obtained at multiple time points, including prior to launch, after the suborbital flight, and on the 4th, 7th, and 10th days of the ISS mission. Additionally, blood samples from three healthy volunteers served as comparative controls. All samples were collected in K2EDTA tubes (10.0 mL, 16 × 100 mm, 18.0 mg, BD Vacutainer®, Cat No: 367525) and stored at −80 °C (NUVE DF 290 E) until further processing. Ethical approvals and participant consents were obtained in accordance with relevant institutional and national guidelines.

### PBMC Isolation

Peripheral blood mononuclear cells (PBMCs) used in this study were isolated according to the experimental protocol previously described by Tastan et al. (2025). Venous blood samples obtained from astronauts and control volunteers were processed following the cellular isolation and culture procedures detailed in that publication **[11**]

### Transcriptome Analysis

Transcriptome analyses were conducted in collaboration with SZA OMICS (Genomic Productions, Istanbul, Turkey) using peripheral blood samples obtained from astronauts who participated in an orbital mission aboard the International Space Station (ISS), an astronaut who completed a suborbital flight (∼100 km), and three healthy volunteer donors. RNA extraction was performed using either the QIAsymphony automated platform (Qiagen, Germany) with the QIAsymphony PAXgene Blood RNA Kit (Catalog No: 762635) or manually with the Qiagen RNeasy Kit, depending on sample characteristics and RNA yield. RNA integrity and concentration were evaluated using NanoDrop, Qubit, and Victor Nivo. Due to suboptimal RIN values (3.9 and 4.2) obtained from some astronaut samples, complementary RNA was derived from corresponding cell cultures to support the analysis. Library preparation was performed using the Illumina Stranded Total RNA Prep, Ligation with Ribo-Zero Plus Kit on the Hamilton NGS STAR platform, followed by sequencing on the Illumina NovaSeq 6000 system in 2×100 bp paired-end format, with an average of 25 million reads per sample. Raw sequencing data were processed with Illumina DRAGEN Bio-IT Platform v3.9.5 and delivered in FASTQ format with Illumina 1.8+ quality encoding. Quality assessment was performed using FastQC, and low-quality and adapter-containing reads were removed using Trimmomatic. Cleaned reads were aligned to the GRCh38 human reference genome using STAR, and transcript expression levels were calculated using TPM normalization. Reference genes for normalization were selected based on pre-analysis results from healthy volunteers.

### ORF Analysis

Open reading frame (ORF) analysis was performed to evaluate the potential protein-coding capacity of the statistically significant LOC genes. The nucleotide sequences of these genes were retrieved in FASTA format from the NCBI GenBank database, and the analyses were conducted using the NCBI ORFfinder tool (https://www.ncbi.nlm.nih.gov/orffinder/). The parameters applied during the analysis included the acceptance of only ‘ATG’ as the start codon, a minimum ORF length threshold of 75 nucleotides, scanning in all six reading frames (+1, +2, +3, −1, −2, 3), and a 5′→3′ reading direction using the standard genetic code (Translation Table 1). All detected ORFs were recorded along with their frame orientation, chain polarity, start and end positions, and the length of the encoded amino acid chains. For each gene, the longest ORF was identified and selected as the representative sequence for further in silico functional evaluations. These findings suggest that LOC genes, which are classically regarded as non-coding, may possess putative translational potential.

### Statistical Analysis

To evaluate the expression levels of LOC genes at five different time points (Pre-launch, Post-suborbital, L+4, L+7, and L+10), the nonparametric Kruskal-Wallis test was applied using GraphPad Prism 8.0.1 software. This test was preferred because normal distribution and homogeneity of variance conditions were not met in the data set (Conover & Iman, 1981). The time points for each gene were treated as independent groups, the data were sorted, the mean rank values were calculated, and the test statistic (H) obtained was compared with the chi-square distribution with k-1 degrees of freedom. As a result of this analysis, significant time-dependent expression changes were found in some LOC genes, while statistical significance could not be obtained in some genes. However, biologically remarkable expression trends were observed in these genes.

## 3. Results

### Conceptual Overview of the MESSAGE Science Mission

The Microgravity Associated Genetics (MESSAGE) Science Mission represents one of Türkiye’s pioneering human space biology initiatives, designed to investigate transcriptional regulation in immune cells under microgravity. As part of the experimental framework, venous blood samples were collected from astronauts seven days before launch to serve as baseline (Earth control) references. Additional samples were obtained onboard the International Space Station (ISS) on mission days 4, 7, and 10 and preserved in the MELFI (Minus Eighty-Degree Laboratory Freezer for ISS) to capture in-flight molecular dynamics. To complement long-duration orbital data, short-term microgravity effects were assessed during the Galaxy-07 suborbital mission. In this phase, blood samples were collected immediately before and after a suborbital flight at ∼100 km altitude, providing an intermediate gravitational condition for comparison. All samples were stored at −80 °C until transcriptomic profiling was performed **(Fig. 1)**. Together, these orbital and suborbital datasets enabled a comprehensive assessment of time-resolved gene expression dynamics across distinct gravitational environments.

**Figure 1:**
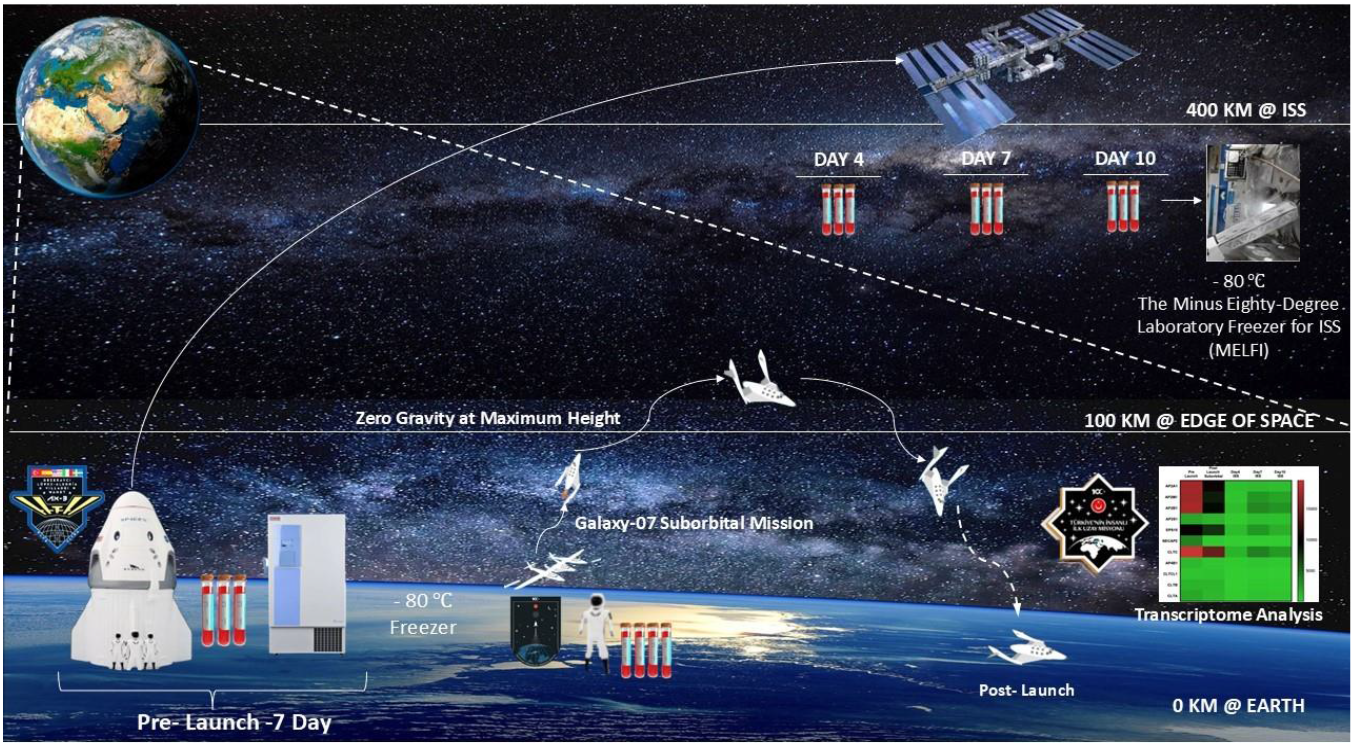
Conceptual framework of the MESSAGE Science Mission. Schematic representation of sample collection and analysis across different mission phases. Blood samples were collected at five time points: Pre-Launch (baseline, Earth control), Post-Launch Suborbital (∼100 km), and Days 4, 7, and 10 aboard the ISS (∼400 km). All samples were cryopreserved at −80 °C, with in-flight samples stored in the ISS MELFI system, and subsequently subjected to transcriptomic profiling upon return to Earth. This design integrates both suborbital and orbital microgravity conditions to capture early and time-dependent transcriptional adaptations.

### Transcriptome Analysis Results

Transcriptome analyses were performed using peripheral blood samples obtained from astronauts aboard the International Space Station (ISS), as well as from three healthy volunteers matched for age, height, and weight. Additionally, samples from a suborbital flight participant who reached an altitude of approximately 100 km were included in the study. The samples were collected at five distinct time points: Pre-Launch, Post-Launch Suborbital, and Days 4, 7, and 10 of the ISS mission. The transcriptional effects of the microgravity environment were evaluated based on this time-course design. The raw RNA sequencing data obtained from these samples were analyzed to assess temporal changes in gene expression. Notably, differences between the Pre-Launch time point and subsequent phases suggest that the transition into space may trigger early transcriptional adaptations. Expression changes observed on ISS Days 4, 7, and 10—conducted in approximately 400 km orbit—highlight molecular alterations occurring at the cellular level under spaceflight conditions. While certain genes exhibited significant upregulation upon exposure to microgravity, others showed suppressed expression profiles. Data from healthy volunteers served as the primary control group for comparative purposes. These data provided a reference framework for evaluating the biological impact of extreme environmental factors such as microgravity. By analyzing astronaut-derived samples across distinct mission phases, the study delineated time-resolved transcriptomic shifts associated with human spaceflight.

### Expression Profiles of LOC Genes Under Microgravity

RNA-Seq analyses revealed dynamic expression profiles of LOC genes across different mission phases, reflecting the transcriptional sensitivity of these uncharacterized elements to microgravity. Six LOC genes showed statistically significant differential expression (p < 0.05, Kruskal–Wallis test). In addition, several other LOC genes, although not reaching statistical significance (p > 0.05), displayed biologically notable expression changes. For example, LOC112268288 demonstrated a pronounced increase after the suborbital flight, whereas LOC101928841 and LOC105373233 exhibited fluctuating patterns across ISS days that, while not statistically significant, suggested potential involvement in microgravity-associated adaptation (**Fig. 2)**. These observations underscore the importance of evaluating both statistically validated results and biologically meaningful, though non-significant, expression shifts when characterizing the molecular response to spaceflight. **Figure 2** presents bar graphs depicting the temporal expression levels of selected LOC genes that did not achieve statistical significance yet displayed noteworthy transcriptional fluctuations. Each bar represents mean normalized expression values (TPM) across biological replicates at Pre-Launch, Post-Launch Suborbital, and ISS Days 4, 7, and 10. Despite the absence of statistical significance, the visual trends highlight subtle but consistent modulations in gene activity, suggesting that these LOC genes may contribute to early or late-stage cellular responses under gravitational stress. The bar graphs serve to emphasize the biological relevance of these temporal changes and provide a foundation for future investigations into the functional potential of LOC transcripts.

**Figure 2:**
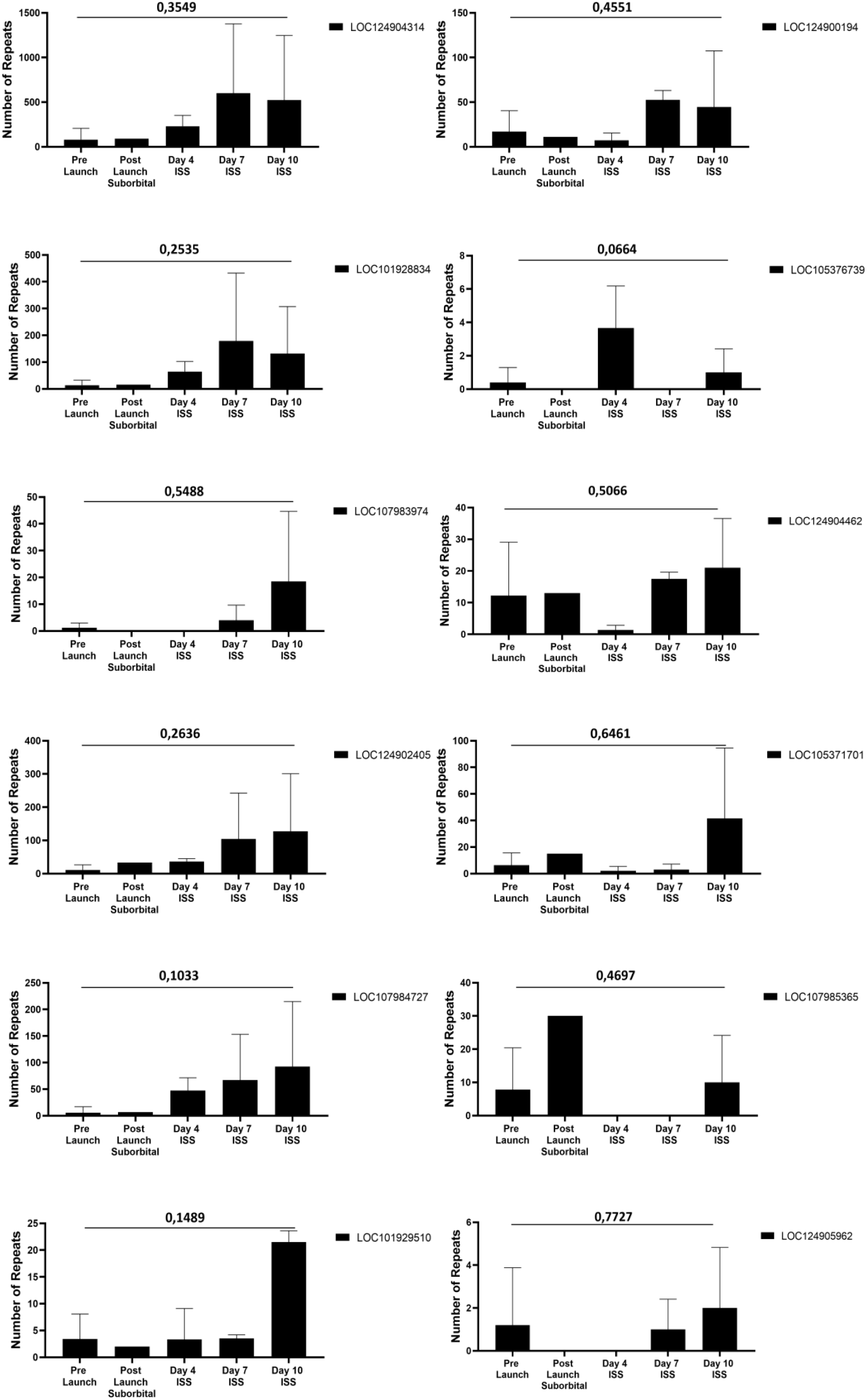
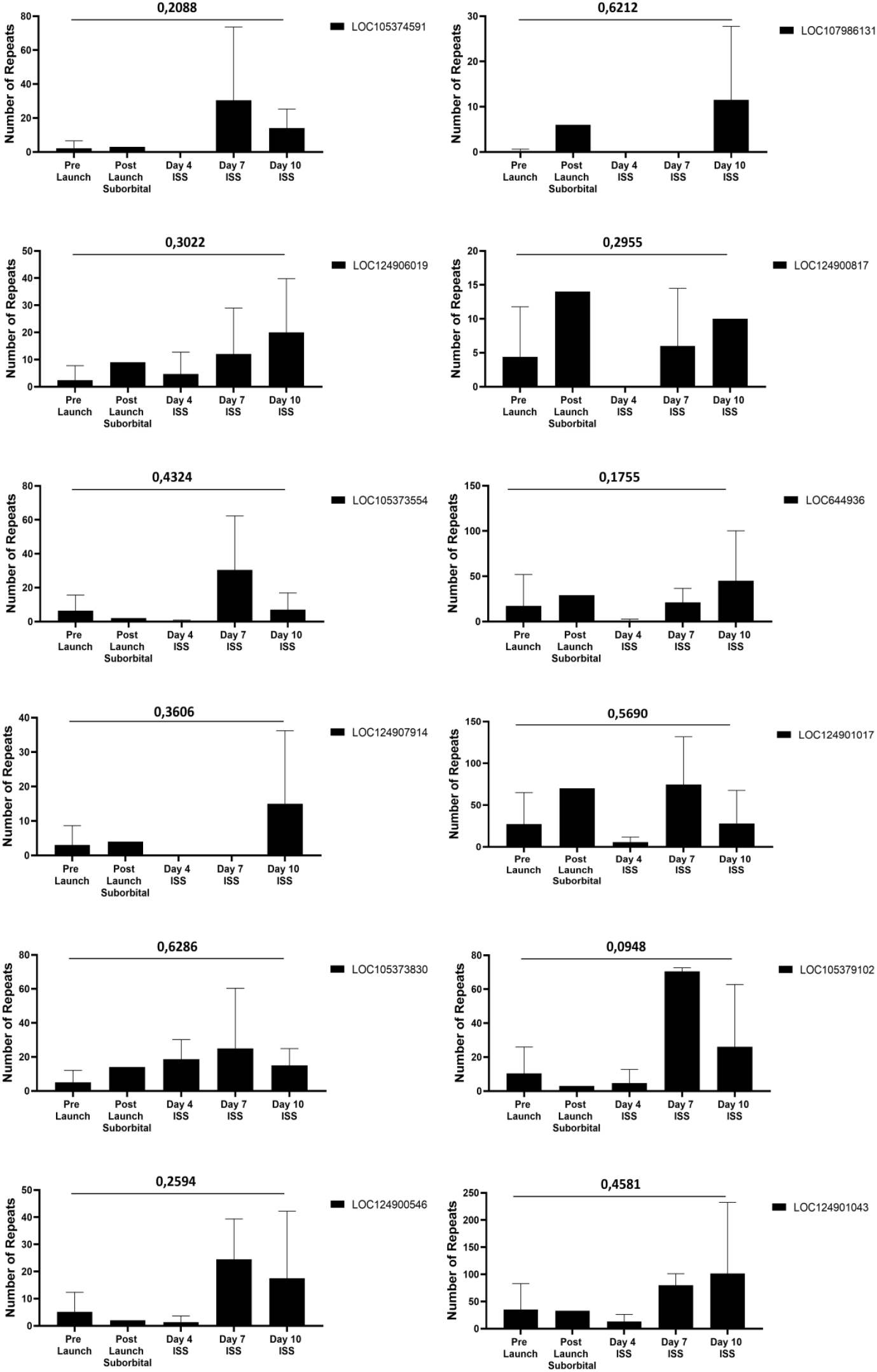
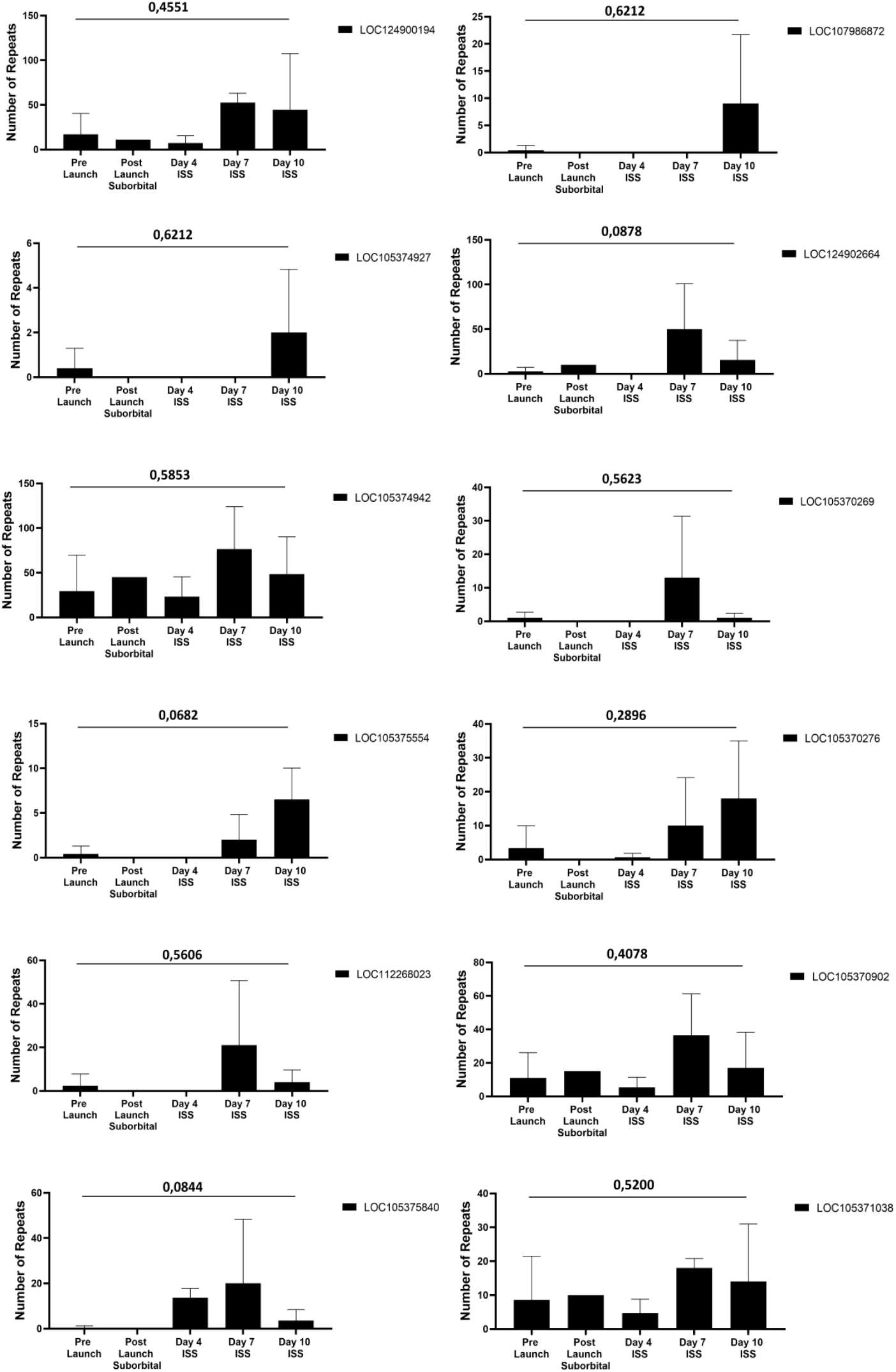
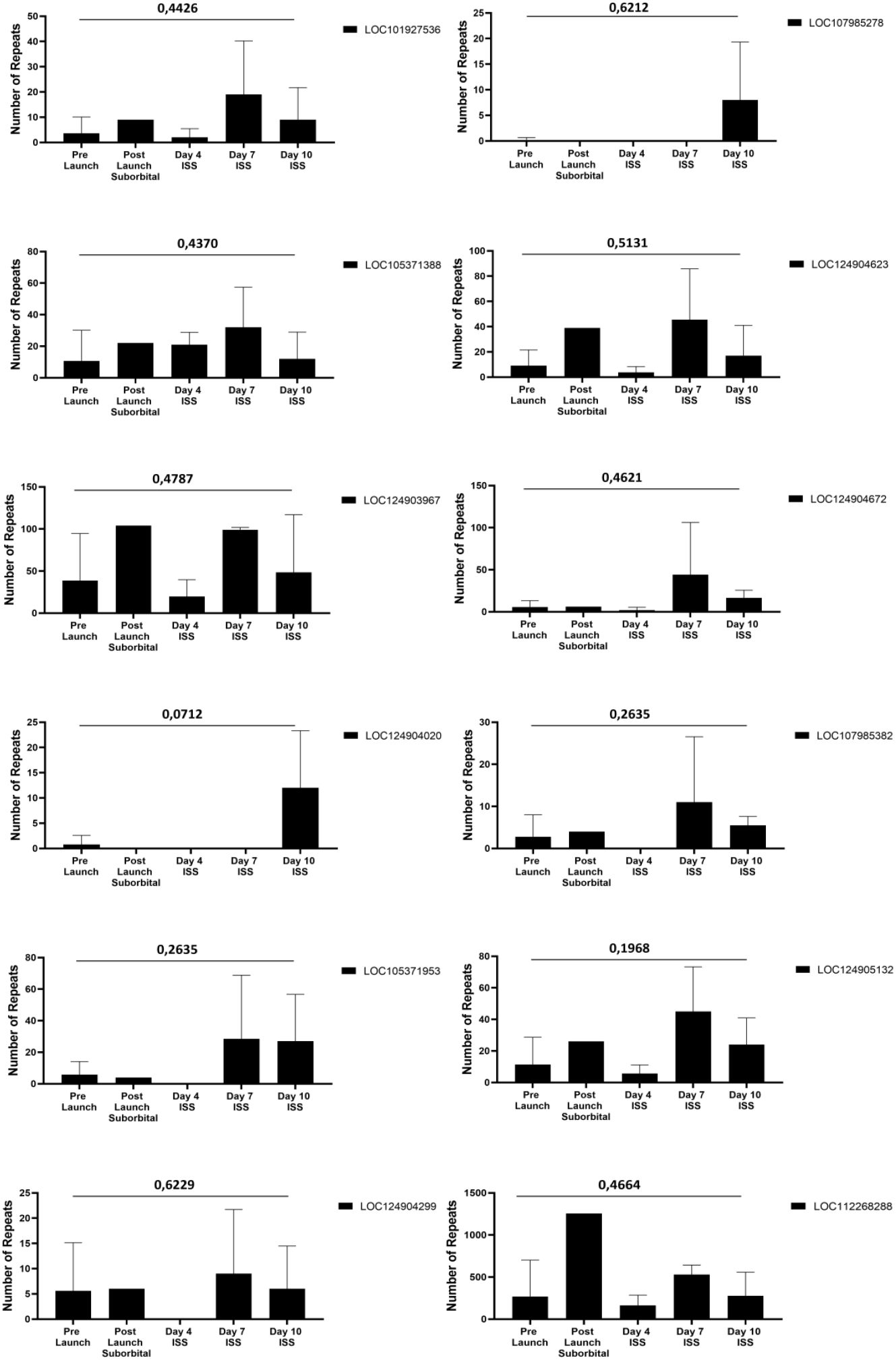
Bar graph representation of LOC genes showing biologically notable but non-significant expression changes under microgravity. Mean normalized expression values (TPM) of selected LOC genes are shown across five mission phases: Pre-Launch, Post-Launch Suborbital (∼100 km), and ISS Days 4, 7, and 10 (∼400 km). Although statistical significance was not achieved (p > 0.05, Kruskal–Wallis test), temporal fluctuations in expression highlight biologically relevant transcriptional responses that may reflect early adaptation mechanisms to microgravity. Error bars represent the standard error of the mean (SEM).

### Heatmap Profiles of LOC Genes Under Microgravity

While bar graphs allow for the visualization of individual gene expression dynamics, they do not fully capture the collective transcriptional patterns across the LOC gene subset. To address this, the data were further consolidated into a heatmap **(Fig. 3)**, which integrates the temporal expression trends of multiple non-significant yet biologically notable LOC genes. This broader visualization highlights subtle but consistent transcriptional modulations across mission phases, providing a more holistic perspective of how these genes may function collectively in the context of microgravity-induced stress adaptation. This consolidated visualization highlights subtle transcriptional modulations across Pre-Launch, Post-Launch Suborbital, and ISS Days 4, 7, and 10, emphasizing temporal changes that may be overlooked when considering each gene in isolation. The heatmap reveals that, while statistical thresholds were not met (p > 0.05, Kruskal–Wallis test), several LOC genes exhibited consistent directional trends. For instance, certain transcripts showed transient upregulation during the suborbital phase, followed by downregulation at later ISS stages, while others demonstrated delayed activation peaking around Day 7. These patterns suggest that even non-significant genes may play auxiliary or regulatory roles in the cellular stress response to microgravity, reflecting early adaptation or individual variability. Importantly, this analysis demonstrates the value of combining statistical outcomes with biological interpretation: genes with non-significant p-values can still exhibit reproducible transcriptional patterns that merit further investigation, particularly in the context of small sample sizes typical of space biology experiments.

**Figure 3:**
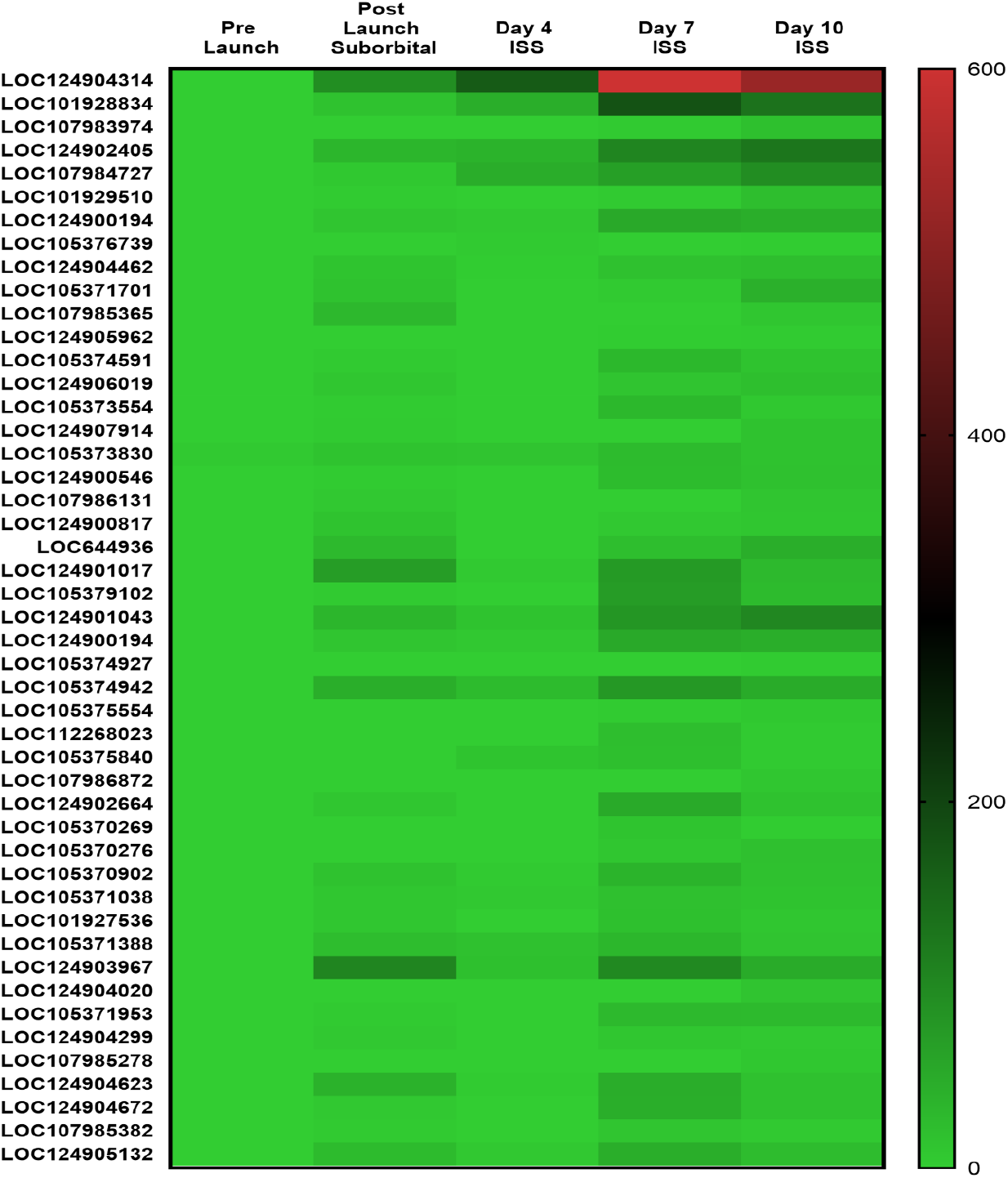
Heatmap of LOC genes displaying non-significant but biologically notable expression profiles under microgravity. Normalized expression values (TPM) of selected LOC genes across Pre-Launch, Post-Launch Suborbital (∼100 km), and ISS Days 4, 7, and 10 (∼400 km) are presented. Although these transcripts did not surpass statistical significance (p > 0.05, Kruskal–Wallis test), the heatmap reveals consistent temporal modulations, including transient activation and delayed expression peaks, suggesting possible roles in early cellular adaptation to microgravity. Color intensity reflects relative expression levels.

### LOC Genes Not Statistically Significant

Although several LOC genes did not achieve statistical significance (p > 0.05, Kruskal–Wallis test), their temporal expression dynamics remain biologically noteworthy. Bar graph and heatmap analyses (Figures 2 and 3) demonstrated that some transcripts consistently fluctuated across mission phases, suggesting sensitivity to gravitational stress despite the absence of strong statistical support. **Figure 4** provides an additional representation of these expression profiles, highlighting genes with observable but variable responses during the suborbital and ISS phases. In particular, some LOC genes displayed transient upregulation immediately after suborbital flight, followed by stabilization at later ISS stages, while others exhibited delayed induction around Day 7 with subsequent return to baseline at Day 10. These expression patterns suggest that certain genes may act as transient responders or modulators of stress adaptation. The lack of statistical significance may be attributed to small sample size, inter-individual genetic variability, and occasional reductions in RNA quality. Nonetheless, documenting these temporal trends is important, as they may inform hypotheses for larger-scale studies and guide the selection of candidate biomarkers in future space biology research.

**Figure 4:**
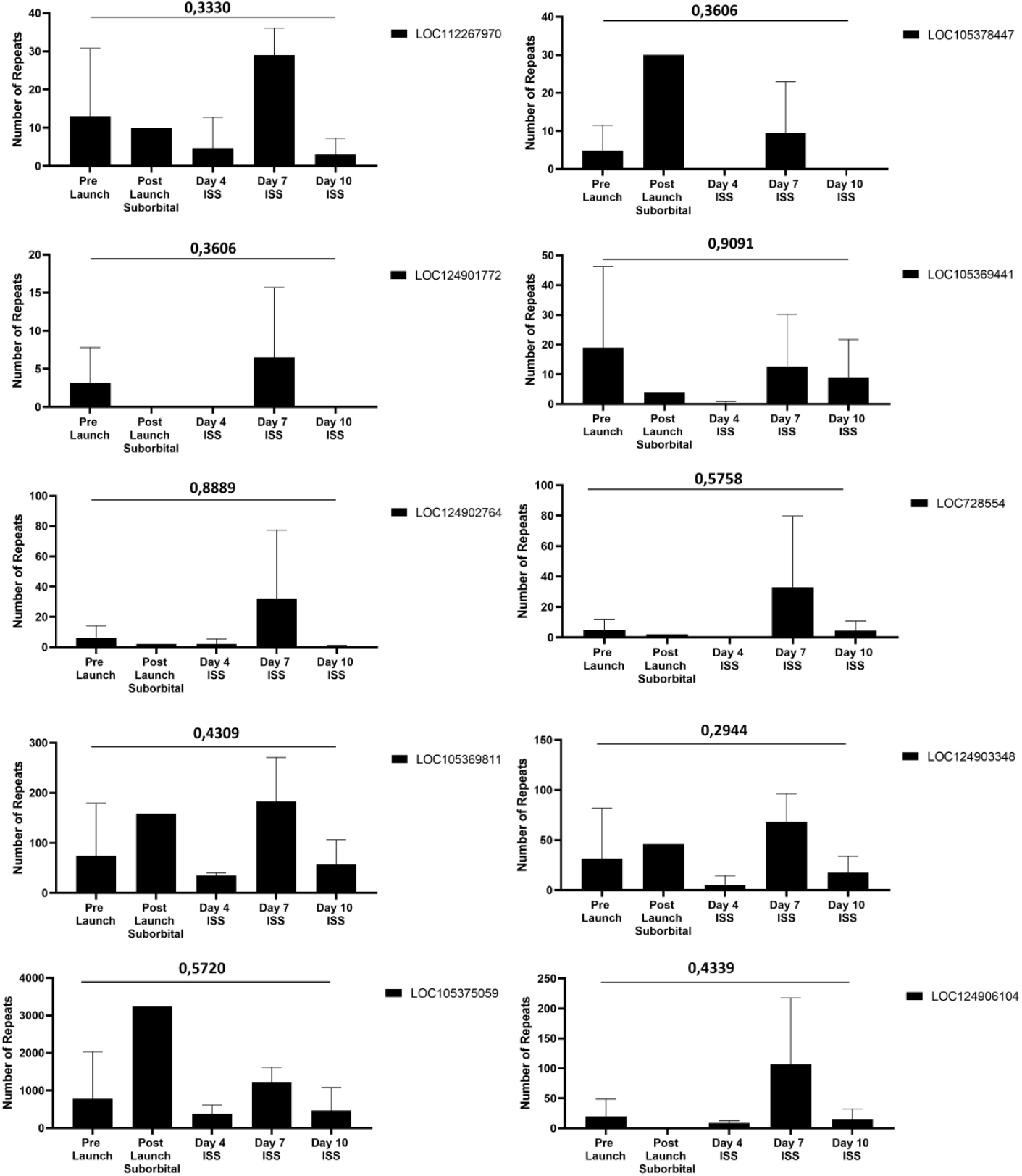
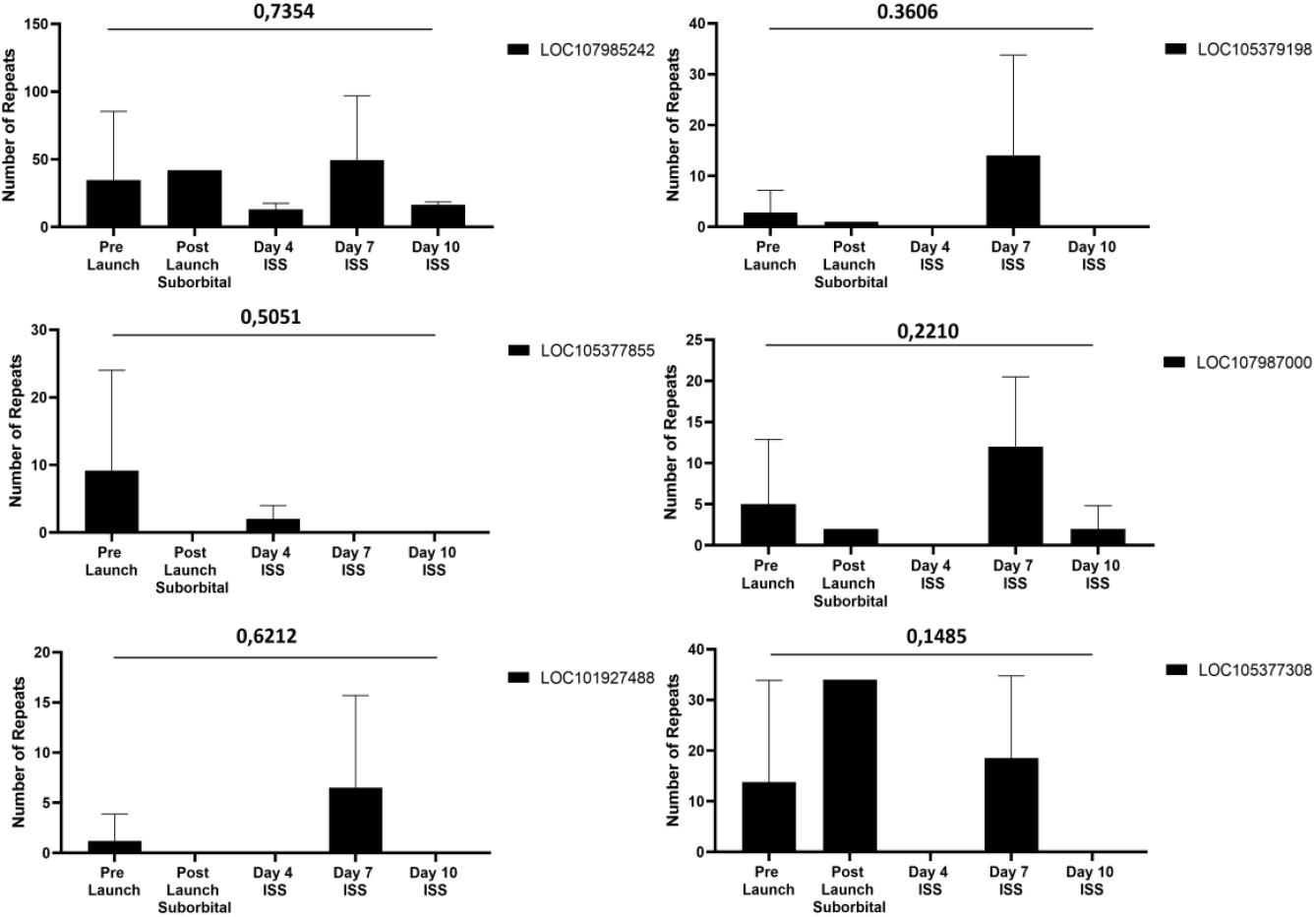
Expression profiles of LOC genes that did not reach statistical significance under microgravity. Line and bar graph visualizations of non-significant LOC genes across five mission phases: Pre-Launch, Post-Launch Suborbital (∼100 km), and ISS Days 4, 7, and 10 (∼400 km). Despite p > 0.05 (Kruskal–Wallis test), several genes exhibited biologically meaningful temporal trends, including transient activation and delayed expression peaks, suggesting potential roles in adaptive responses to gravitational stress.

### Statistically Significant LOC Genes Under Microgravity

In contrast to the non-significant but biologically notable transcripts, six LOC genes demonstrated statistically significant expression changes across mission phases (p<0.05, Kruskal–Wallis test). Among these, LOC117779438 and LOC102724934 displayed particularly dynamic transcriptional profiles. LOC117779438 exhibited a progressive and mission-sensitive increase, with upregulation becoming most prominent after the suborbital flight and peaking at ISS Day 10. This trend suggests a sustained and cumulative response to gravitational stress. Conversely, LOC102724934 demonstrated a more complex temporal pattern. Expression levels rose following the suborbital flight and ISS Day 4, dropped sharply to near-baseline levels on Day 7, and reappeared at Day 10. Such a discontinuous trajectory implies that this gene may function as a timing-sensitive responder, activated under specific stress windows before returning to homeostatic levels. These findings highlight the diversity of transcriptional responses among LOC genes, with some showing continuous activation and others demonstrating oscillatory or transient dynamics. Together, these results underscore the potential of LOC transcripts to act as molecular indicators of spaceflight adaptation, reflecting both immediate and delayed regulatory mechanisms in response to microgravity.

### Genes Activated in Suborbital and Space but Homeostasis Over Time

Some LOC genes displayed statistically significant expression increases during the suborbital and early ISS mission phases, followed by a return to baseline or decreased expression at later time points. Notably, LOC105370844 and LOC105372255 exemplify this transient activation profile. LOC105370844 was significantly upregulated after the suborbital flight and again at ISS Day 7, but its expression declined sharply by Day 10, suggesting an early stress-induced activation followed by adaptation. Similarly, LOC105372255 peaked at ISS Day 4 before progressively declining in subsequent days, indicating an early but short-lived transcriptional response. These results suggest that certain LOC genes act as transient responders to microgravity, initiating stress adaptation mechanisms during the initial phases of exposure but returning to equilibrium as homeostatic processes stabilize cellular function. Such patterns highlight the importance of temporal resolution in space biology studies, as early activation could be overlooked in studies sampling only at later time points **(Fig. 6)**.

### LOC Genes with Delayed Activation During Spaceflight

Another subset of LOC genes demonstrated late-response profiles, characterized by little to no expression during early mission stages but significant induction at later ISS phases. LOC124905103 and LOC124900480 showed minimal expression during the suborbital and early ISS phases but were strongly upregulated at ISS Day 7. Interestingly, both genes subsequently declined toward homeostatic levels by Day 10, indicating a transient but delayed activation pattern. These delayed-response genes may reflect molecular processes that are triggered after prolonged exposure to microgravity, such as cumulative stress signaling, transcriptional reorganization, or secondary regulatory feedback loops. Their expression profiles suggest that LOC genes may participate in temporally distinct adaptation mechanisms, ranging from early transient responders **(Fig. 6)** to delayed activation and resolution dynamics **(Fig. 7)**.

### Integrated Heatmap of Statistically Significant LOC Genes

To consolidate the temporal expression dynamics of the six statistically significant LOC genes, a heatmap analysis was performed (Figure 8). While Figures 5–7 illustrate individual response profiles—ranging from continuous activation (LOC117779438), to transient responders (LOC105370844 and LOC105372255), and delayed activators (LOC124905103 and LOC124900480)—the heatmap provides a comprehensive overview of their collective regulation under microgravity. This visualization highlights distinct but complementary transcriptional patterns: some genes exhibited progressive upregulation across mission phases, while others showed transient or oscillatory expression profiles. Together, these trajectories reflect the heterogeneity of molecular responses to gravitational stress and underscore the role of LOC genes as a diverse regulatory network rather than isolated responders. Importantly, the clustering observed within the heatmap suggests functional convergence among certain genes, supporting the hypothesis that subsets of LOC transcripts may operate in coordinated adaptation pathways during spaceflight.

**Figure 5:**
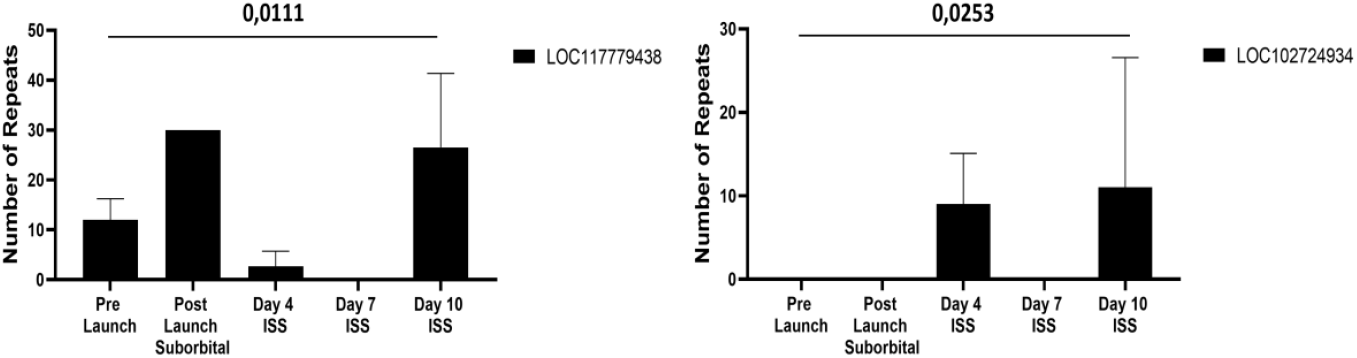
Statistically significant LOC genes showing dynamic expression under microgravity. Bar graphs of LOC117779438 and LOC102724934 expression levels across mission phases: Pre-Launch, Post-Launch Suborbital (∼100 km), and ISS Days 4, 7, and 10 (∼400 km). LOC117779438 exhibited continuous upregulation with a strong increase at later ISS stages, while LOC102724934 showed a fluctuating pattern with intermittent activation. Both genes surpassed the statistical threshold (p < 0.05, Kruskal–Wallis test), demonstrating distinct transcriptional adaptations to microgravity. Error bars represent the standard error of the mean (SEM).

**Figure 6:**
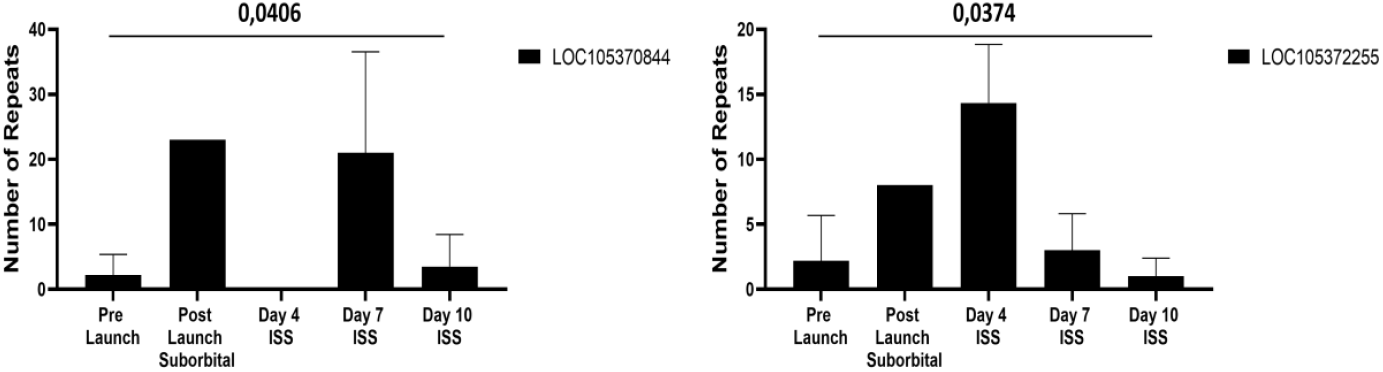
LOC genes are transiently activated under microgravity but return to homeostasis. Expression dynamics of LOC105370844 and LOC105372255 across Pre-Launch, Post-Launch Suborbital (∼100 km), and ISS Days 4, 7, and 10 (∼400 km). Both genes showed significant activation during early mission phases, followed by decreased expression and stabilization toward baseline levels by Day 10. Statistical significance was confirmed (p < 0.05, Kruskal–Wallis test). Error bars represent the standard error of the mean (SEM).

**Figure 7:**
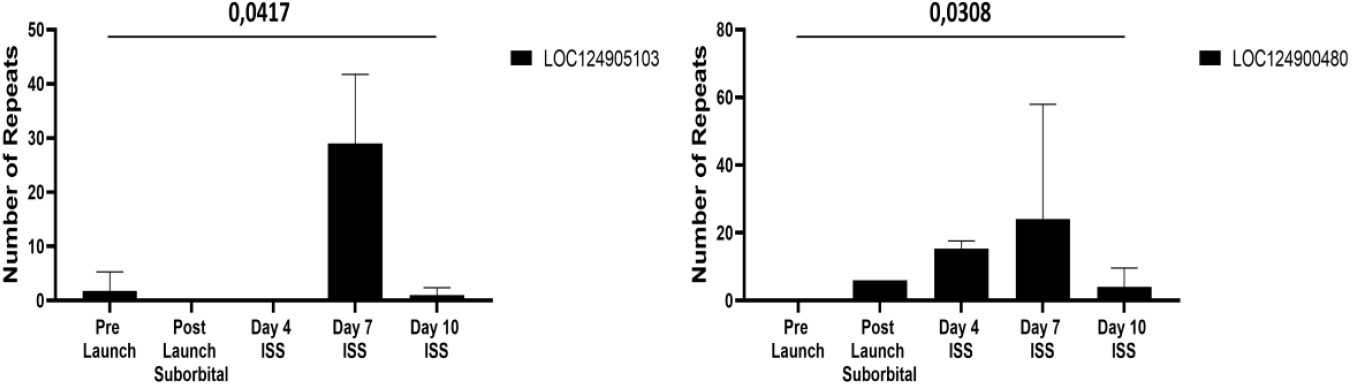
LOC genes showing delayed activation during ISS missions. Expression dynamics of LOC124905103 and LOC124900480 across Pre-Launch, Post-Launch Suborbital (∼100 km), and ISS Days 4, 7, and 10 (∼400 km). Both genes showed minimal activity during early phases but demonstrated significant upregulation at ISS Day 7, followed by a return toward baseline at Day 10. Statistical significance was confirmed (p < 0.05, Kruskal–Wallis test). Error bars represent the standard error of the mean (SEM).

**Figure 8:**
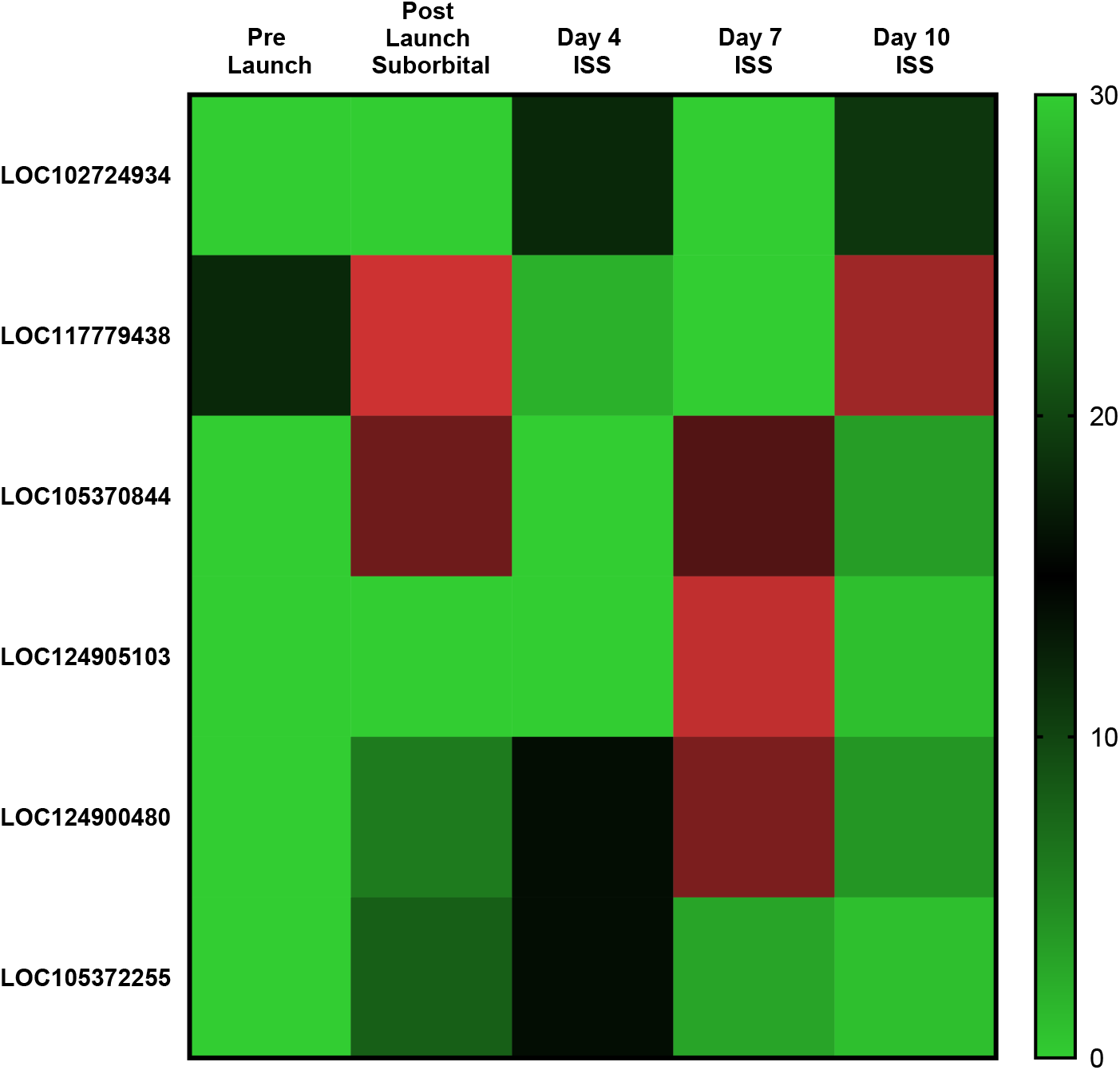
Heatmap of statistically significant LOC genes under microgravity. Temporal expression patterns of six LOC genes (p < 0.05, Kruskal–Wallis test) across five mission phases: Pre-Launch, Post-Launch Suborbital (∼100 km), and ISS Days 4, 7, and 10 (∼400 km). The heatmap integrates diverse transcriptional responses— including progressive activation, transient induction, and delayed expression—into a collective profile of microgravity-associated adaptation. Color intensity reflects relative normalized expression values (TPM).

### Phylogenetic Proximity Between Significant LOC Genes

To explore potential evolutionary relationships among the six statistically significant LOC genes, multiple sequence alignment (MSA) and phylogenetic analysis were conducted. The resulting tree revealed notable clustering patterns that paralleled both expression dynamics and ORF structures. Specifically, LOC124905103 and LOC124900480 were positioned as the closest pair, consistent with their highly similar temporal activation profiles and the presence of identical ORF33 structures. This strong sequence similarity suggests a possible shared evolutionary origin and functional convergence in microgravity response mechanisms. Other genes exhibited greater sequence divergence. For example, LOC102724934 was located at a distal branch of the phylogenetic tree, reflecting its unique fluctuating expression pattern and shorter ORF5 structure. Meanwhile, LOC117779438, LOC105370844, and LOC105372255 formed intermediate subclusters, showing moderate sequence similarity but distinct transcriptional trajectories **(Fig. 9)**. These findings indicate that statistically significant LOC genes not only differ in their temporal responses to microgravity but also in their evolutionary context. Such diversity suggests that microgravity-induced adaptation may recruit a broad spectrum of uncharacterized genomic elements, ranging from closely related transcripts with conserved structures to highly divergent loci with specialized functions.

**Figure 9:**
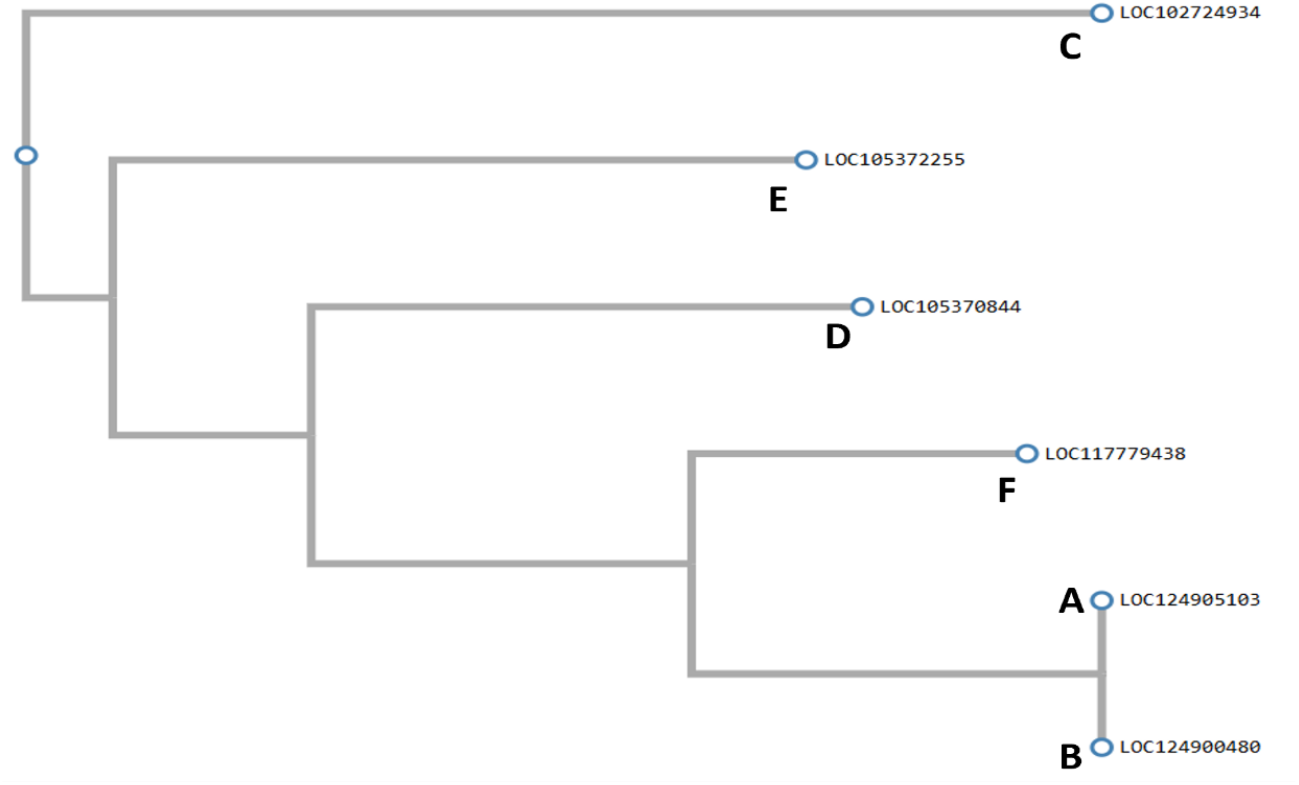
Phylogenetic tree of statistically significant LOC genes. Multiple sequence alignment of six statistically significant LOC genes was performed using Clustal Omega, and a phylogenetic tree was constructed to evaluate evolutionary relationships. LOC124905103 and LOC124900480 clustered closely, consistent with their shared ORF33 structure and similar expression dynamics, while LOC102724934 was placed at a distant branch, reflecting greater sequence and functional divergence. Other genes (LOC117779438, LOC105370844, LOC105372255) were grouped into intermediate subclusters, highlighting both sequence variability and transcriptional diversity among the genes responsive to microgravity.

### Expressional Dynamics Supported by ORF Potential

To further evaluate the functional relevance of statistically significant LOC genes, open reading frame (ORF) analyses were performed. These analyses revealed that several of the transcripts, traditionally annotated as non-coding, possess coding potential. Notably, LOC124905103 and LOC124900480 contained an identical ORF33 structure (882 nt / 293 aa) in the +2-reading frame, a finding consistent with both their close phylogenetic proximity **(Fig. 10)** and their synchronized expression patterns under microgravity. The structural and transcriptional overlap between these two genes suggests the possibility of shared translational products or conserved functional roles in stress adaptation. In contrast, LOC102724934 carried a shorter ORF5 (243 nt / 80 aa) in the +2 frame, reflecting a divergent translational potential that parallels its unique oscillatory expression dynamics. Similarly, significant ORFs were identified in LOC105370844, LOC105372255, and LOC117779438, with the latter harboring an ORF in the −3 frame of the negative strand. This unusual orientation indicates strand-specific regulation and highlights the complexity of translational activity in uncharacterized genomic regions. Taken together, these results support the hypothesis that a subset of LOC genes, despite their classification as “uncharacterized” or “non-coding,” may encode functional peptides under extreme conditions such as microgravity. The convergence of expression dynamics, evolutionary proximity, and ORF structures underscores the need to reconsider the biological significance of LOC transcripts in both spaceflight and terrestrial biology. When these findings are evaluated together, it is concluded that LOC genes, which are classically classified as non-coding, contain structures that may carry translational potential under extreme conditions such as microgravity (Bazzini et al., 2014).

**Figure 10:**
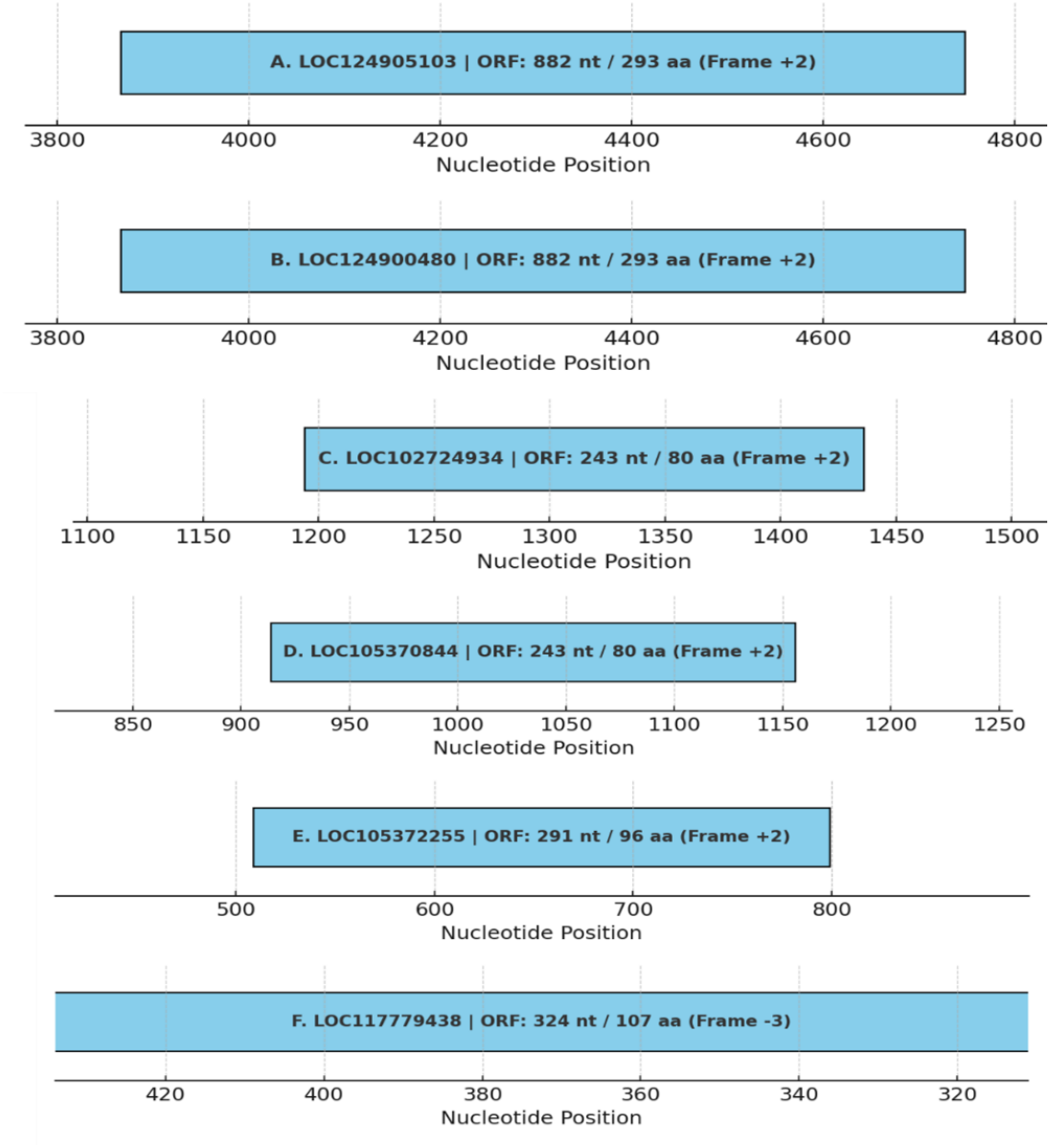
Predicted open reading frames (ORFs) of statistically significant LOC genes. Schematic representation of the longest ORFs identified in six significant LOC genes using NCBI ORFfinder. Blue bars indicate ORF position, length (nt/aa), and translation frame. LOC124905103 and LOC124900480 shared an identical ORF33 (882 nt / 293 aa, Frame +2), consistent with their phylogenetic similarity and expression overlap. LOC102724934 harbored a shorter ORF5 (243 nt / 80 aa, Frame +2), while LOC117779438 featured an ORF in the −3 frame on the negative strand, suggesting strand-specific regulation. These findings demonstrate that LOC genes previously annotated as non-coding may possess translational potential under microgravity conditions.

## 4. Discussion

Microgravity represents a unique and extreme environmental condition for biological systems that have evolved under Earth’s constant gravitational field. Its effects extend beyond well-characterized protein-coding genes, influencing unannotated or poorly defined genomic regions that are often excluded from conventional analyses. In this study, we systematically examined the behavior of LOC (Locus) genes—transcripts frequently annotated as long non-coding RNAs but lacking full functional characterization—and demonstrated that several exhibit significant transcriptional modulations under spaceflight conditions. Our findings highlight three complementary layers of evidence supporting the potential biological relevance of LOC genes in microgravity: (i) expression dynamics, where both statistically significant and biologically notable but non-significant genes displayed time-dependent regulation; (ii) phylogenetic relationships, which revealed evolutionary clustering among transcripts with similar expression and ORF features; and (iii) ORF potential, which showed that a subset of LOC genes harbors open reading frames suggestive of translational capacity. Together, these results challenge the assumption that LOC genes are functionally inert and instead position them as candidate regulators or even coding elements involved in cellular adaptation to gravitational stress. By integrating transcriptomic, phylogenetic, and ORF analyses, this study expands the molecular landscape of space biology beyond canonical gene pathways and into the so-called “dark genome.” These insights suggest that LOC genes may serve as sensitive biomarkers of microgravity-induced stress and provide a foundation for future investigations into their functional and therapeutic relevance.

Microgravity represents an unusual environmental condition for biological systems that have evolved under Earth’s gravitational forces. This condition can lead to profound molecular alterations, particularly in gene expression profiles [9, 3]. While the differential expression of LOC genes under microgravity provides compelling evidence of their potential functional roles, the biological interpretation requires careful consideration. Some transcripts, such as LOC117779438, showed continuous activation across mission phases, suggesting a sustained role in cellular stress adaptation. Others, including LOC105370844 and LOC105372255, demonstrated transient activation followed by a return to baseline, consistent with early-response genes that initiate but do not maintain adaptation pathways. In contrast, LOC124905103 and LOC124900480 exhibited delayed upregulation, peaking at ISS Day 7, which may indicate involvement in secondary or downstream regulatory mechanisms triggered after prolonged exposure. These temporally distinct expression profiles emphasize that LOC genes may participate in multiple phases of adaptation, from immediate stress responses to long-term reorganization of cellular processes. While most previous studies have focused on well-characterized, protein-coding genes, regions of the genome with unknown or poorly defined functions have largely been overlooked in the context of microgravity. This study addressed this gap by focusing on genes annotated with the LOC (Locus) identifier—transcripts often lacking full annotation, unclear in function, and frequently classified as long non-coding RNAs (lncRNAs) [10, 16]. Through transcriptomic analysis, six LOC genes exhibiting statistically significant differential expression were identified using the non-parametric Kruskal-Wallis test, which was chosen due to the limited sample size and non-normal distribution of the data. This methodological decision aligns with recommended practices for small-scale biological studies [15]. These findings suggest that microgravity affects not only classical genetic pathways but also understudied genomic elements. Notably, LOC124900480 and LOC124905103 genes showed significant upregulation on Day 7 of the spaceflight mission. This time point is widely regarded in space biology literature as a critical phase of early cellular reorganization, reflecting the onset of molecular adaptation to microgravity [8]. These results indicate that even short-term space missions can elicit significant transcriptional responses.

The identification of open reading frames (ORFs) within these transcripts further strengthens the case for their biological relevance. The discovery of identical ORF33 structures in LOC124905103 and LOC124900480, combined with their phylogenetic proximity, suggests evolutionary conservation and a potential shared translational role. Meanwhile, the strand-specific ORF identified in LOC117779438 highlights the structural diversity of these genes and raises the possibility of unconventional regulatory mechanisms, such as reverse-strand transcription under stress. These findings align with recent evidence that a subset of long non-coding RNAs may encode functional peptides when cells are exposed to environmental challenges, including oxidative stress and metabolic perturbations. Additionally, some LOC genes exhibited biologically relevant expression patterns despite lacking statistical significance. For example, LOC112268288 demonstrated an apparent increase in expression following suborbital flight; however, high sample variability prevented this from reaching statistical significance. Such cases may reflect individual-specific genetic responses or late-stage adaptation processes. It is increasingly acknowledged in the literature that genes with biologically meaningful trends, even without statistical significance, should be carefully considered [12].To better assess the functional potential of these transcripts, open reading frame (ORF) analyses were conducted using the NCBI ORFfinder tool. These analyses identified ORFs that began with an ATG start codon and ended with a stop codon.

LOC124900480 and LOC124905103 shared an identical ORF33 structure (882 nt / 293 aa) in the same reading frame (+2), demonstrating structural and transcriptional similarity. Phylogenetic analysis further revealed that these two genes clustered closely, suggesting a potential shared molecular function. In contrast, LOC102724934 featured a shorter ORF5 (243 nt / 80 aa) located in a different reading frame, highlighting that translational potential cannot be explained solely by sequence similarity. These results are consistent with studies suggesting that certain lncRNAs may encode proteins under stress conditions [13].This study is not without limitations. The small sample size and variable RNA quality in some samples may have limited statistical power. However, these limitations were transparently reported, ensuring methodological clarity and laying the groundwork for future hypothesis-driven research [17].

This study systematically evaluated the expression patterns and potential functional capacities of LOC (Locus) genes under microgravity conditions, using transcriptomic and ORF-based approaches. The results revealed that several LOC genes, traditionally considered non-coding or functionally unknown, exhibited dynamic expression profiles in response to spaceflight-induced stress. Notably, six LOC genes demonstrated statistically significant changes in expression, and ORF analysis further supported their potential translational activity. These findings suggest that LOC genes, often overlooked in genomic research, may act as sensitive molecular responders under extreme environmental conditions and could represent novel regulatory or coding elements of the human genome. Despite these insights, several limitations must be acknowledged. The small number of astronaut samples inherently reduced statistical power and increased variability, which may explain why some biologically relevant expression trends did not achieve statistical significance. In addition, RNA quality from certain spaceflight samples was suboptimal, necessitating complementary analysis from cultured cells. While these methodological adjustments ensured robust data generation, they may have introduced subtle biases into expression estimates. Finally, inter-individual genetic variability likely contributed to expression heterogeneity, underscoring the need for larger cohorts and replication across multiple missions. Nevertheless, these limitations are common in space biology studies, where sample availability is restricted, and the transparent reporting of such challenges strengthens the reliability of the findings.

In conclusion, this study provides novel evidence that LOC genes—traditionally regarded as uncharacterized or non-coding—exhibit dynamic and functionally relevant transcriptional changes under microgravity. By integrating temporal expression profiles, phylogenetic analysis, and ORF prediction, we demonstrate that these transcripts may act as sensitive molecular responders to gravitational stress, with potential roles that extend beyond regulatory activity to include translational functions. The identification of conserved ORF structures, particularly in LOC124905103 and LOC124900480, reinforces the possibility that these genes encode peptides with adaptive significance in extreme environments. The broader implications of these findings are twofold. First, they expand the scope of space biology beyond canonical protein-coding genes, underscoring the need to investigate the “dark genome” as a reservoir of molecular elements that contribute to human adaptation in space. Second, they point to LOC genes as potential biomarkers of physiological stress during space missions, offering opportunities for early detection of molecular changes that precede clinical manifestations. As spaceflight durations increase and human presence in extraterrestrial environments becomes more routine, such biomarkers play a critical role in monitoring astronaut health and developing targeted countermeasures. Future studies should build on this work by validating LOC gene functions in larger astronaut cohorts, simulating microgravity in ground-based models, and exploring the translational potential of their predicted peptides. Functional characterization through proteomic approaches and gene-editing experiments will be essential to clarify whether these transcripts encode biologically active proteins or operate through non-coding regulatory mechanisms. Ultimately, this research highlights LOC genes as previously overlooked but potentially crucial components of the molecular response to microgravity, providing a foundation for both fundamental genomic discovery and the development of spaceflight-relevant therapeutic strategies.

## Acronyms/Abbreviations

MESSAGE: (Microgravity Associated Genetics) Science Mission
(ISS: International Space Station
(PBMC): Peripheral Blood Mononuclear Cells
(TÜBİTAK): Scientific and Technological Research Council of Türkiye
(TUA): Turkish Space Agency
(RNA-Seq): RNA Sequencing
(TRGENMER): Transgenic Cell Technologies and Epigenetic Application and Research Center

## Funding Acknowledgement

This study was supported by the Scientific and Technological Research Council of Türkiye (TÜBİTAK) and the Turkish Space Agency (TUA) through the Turkish Astronaut and Scientific Mission (TABM) Project (1007-KAMAG-121L002 / Agreement No: TABM-HZT-23-25). We would like to thank TÜBİTAK UZAY and TUA for their support and declare that none of the opinions and findings contained in this publication are the official views of TÜBİTAK and TUA. The research process was initially launched through two TÜBİTAK 2209-A Projects (Application Nos: 1919B012112510 and 1919B012208729). This was followed by a Scientific Research Project at Üsküdar University (Project No: ÜÜBAP-YP-2021-016), which focused on the development of a device capable of simulating microgravity conditions.

## Author Contributions

O.D.: Data curation, Formal analysis, Visualization, Investigation, Writing – original draft, Writing – review & editing. E.C.: Data curation, Formal analysis, Investigation. B.Y.: Software, Methodology, Validation. K.K.: Software, Data curation, Formal analysis. M.R.S.: Visualization. B.A.: Visualization, Writing – review & editing. C.T.: Conceptualization, Methodology, Supervision, Project administration, Writing – original draft, Writing – review & editing.

## Acknowledgements

The experience and infrastructure gained through these efforts provided the foundation for participation in the MESSAGE Science Mission. We gratefully acknowledge the support of TUA, TÜBİTAK, and Üsküdar University throughout this process. The opinions and findings expressed in this publication are solely those of the authors and do not necessarily represent the official views of the supporting institutions. Finally, we extend our sincere thanks to all research colleagues, data analysts, and technical staff whose contributions were essential to the success of the MESSAGE Science Mission.

## Notes

### Competing Interest Statement

The authors have declared no competing interest.

